# Groping in the fog: soaring migrants exhibit wider scatter in flight directions and respond differently to wind under low visibility conditions

**DOI:** 10.1101/2021.07.22.453357

**Authors:** Paolo Becciu, Michele Panuccio, Giacomo Dell’Omo, Nir Sapir

## Abstract

Atmospheric conditions are known to affect flight propensity, behaviour during flight, and migration route in birds. Yet, the effects of fog have only rarely been studied although they could disrupt orientation and hamper flight. Fog could limit the visibility of migrating birds such that they might not be able to detect landmarks that guide them during their journey. Soaring migrants modulate their flight speed and direction in relation to the wind vector to optimize the cost of transport. Consequently, landmark-based orientation, as well as adjustments of flight speed and direction in relation to wind conditions, could be jeopardized when flying in fog. Using a radar system operated in a migration bottleneck (Strait of Messina, Italy), we studied the behaviour of soaring birds under variable wind and fog conditions over two consecutive springs (2016 and 2017), discovering that migrating birds exhibited a wider scatter of flight directions and responded differently to wind under fog conditions. Birds flying through fog deviated more from the mean migration direction and increased their speed with increasing crosswinds. In addition, airspeed and groundspeed increased in the direction of the crosswind, causing the individuals to drift laterally. Our findings represent the first quantitative empirical evidence of flight behaviour changes when birds migrate through fog and explain why low visibility conditions could risk their migration journey.

## Introduction

Atmospheric conditions are known to affect flight propensity, behaviour during flight, and migration route in birds [1–5]. Migration routes can be several thousand kilometres long, across many different ecosystems, landscapes, and climatic regions. Migratory birds are adapted to fly under different weather conditions, but in general, they prefer to fly in favourable atmospheric conditions. For instance, massive nocturnal migration in North America is triggered by mild temperatures [6], and flight over ecological barriers is facilitated by tailwinds or weak headwinds and is avoided under strong headwinds [7–11].Fog and low clouds may lower the visibility of landmarks and hamper orientation [5, 12–14]. Fog usually occurs in calm weather conditions (e.g., weak or no winds) near ground level, and its presence might be associated with otherwise good conditions for migration [15]. Although birds may benefit from such weather, the low visibility associated with fog may cause disorientation and consequently avoidance of flight [16, 17]. Bird flight behavior and movement paths could consequently be affected by fog. For example, tracks of Sandhill Cranes (*Antigone canadensis*) recorded on a foggy day with a marine radar were more tortuous and circuitous than on days with good visibility [18]. If the fog extends over a large area, birds could find themselves tens or even hundreds of kilometres away from their intended migratory routes and may become exhausted, as recorded for a flock of Turkey Vultures (*Cathartes aura*) flying over a fog-covered sea where the birds eventually alighted on a boat [19]. Fog may even cause mass-mortality events of migrating birds [20] and may postpone their departure from stopover sites as they wait for better weather conditions. In some cases, birds may even undertake reverse migration when visibility is poor [12, 16, 21, 22]. Indeed, fog was found to delay the arrival of birds at an offshore island in California [23]. These mostly anecdotical findings highlight the diffculty in studying how fog influences bird behaviour. Yet, more comprehensive studies regarding the effects of fog on wildlife can be undertaken in areas where fog prevails over long time periods. For example, Panuccio et al. (2019)[17], who used radar to study soaring migrants in a migratory bottleneck in Southern Italy, documented a substantial decrease in migration intensity under foggy conditions, likely indicating avoidance behaviour. Wind conditions can hamper or facilitate bird migration [5, 7, 24–27]. Specifically, due to the benefits of flying with favourable wind, tailwind assistance could lead to a decision to depart, resulting in a high migration traffic rate [28–30]. To reduce their flight cost and migration duration [31–33] flying migrants should continuously adjust their flight behaviour in response to changes in wind speed and direction as predicted by the optimal migration flight theory [34–36]. Consequently, their air-and groundspeed should change in a predictable manner [33]. Several studies have shown that both flapping and soaring birds reduce their airspeed under tailwinds and increase it under headwinds [34, 35, 37] and that the migrants’ ground speed is also affected by tailwinds, headwind and crosswinds [33, 38–41]. When gliding, soaring migrants reduce airspeed in tailwinds, which effectively increases distance covered [38]. This thereby reduces the chances of grounding and the need to switch to energetically costly flapping flight [42, 43]. The aim of this study was to examine the effects of fog on orientation and response to wind in soaring migrants tracked by a marine radar over a migration bottleneck, The study was carried out at the Strait of Messina in Southern Italy, where Honey Buzzards (*Pernis apivorous*) comprise approximately 95% of the tracked migrants [33, 44]. We analysed the distribution of flight directions and the effects of wind conditions (speed and direction) on buzzard ground-and airspeed when they fly in fog versus clear conditions. We predicted a larger scatter of flight directions and different response to wind in fog, although to the best of our knowledge, no theory exists for flight behaviour in fog. In clear air, bird airspeed and flight direction are expected to change in predictable ways in relation to wind speed and direction to optimize the cost of transport [34, 45]. Whether birds will behave optimally in fog is not known and may depend on the information available to them under these conditions and no previous study has quantitatively estimated how flight may change under these conditions. If they rely on landmarks to calibrate their seasonal direction of flight and speed, and these landmarks are not visible in fog, the birds may risk flying off track and may not notice potential collision hazards simply due to lack of information required to assess their position and movement progress.

## Methods

### Study area and data collection

We collected the data during the spring migration of 2016 (24^th^ March – 7^th^ May) and 2017 (22^nd^ March – 23^rd^ May) near the edge of a flat highland frequently exposed to fog [17] in the Aspromonte Mountains, Calabria (38°23’ N, 15°79’ E – 1030 m a.s.l.), about 7 km inland from the Strait of Messina in southern Italy (Fig. 1). Fog and low clouds are generated in this area because humid air is trapped between the coast and the highland [17]. We collected data using a 12 kW, X-band (9.1 GHz) marine radar rotating at 38 RPM with a 2.1-m antenna that was set horizontally with a 22°vertical beam. The radar covered a sector of 240°(it was blanked towards the observers for the remaining angle) and its detection radius was about 2 km, orienting towards the prevalent direction of the incoming migrants (a compass direction of 215°). Its horizontally rotating antenna enabled computing the direction and speed of the migrating birds based on the detection of their echoes and the reconstruction of their trajectories. Notably, we could not compute the altitude of the echoes. The radar was positioned in an area that is well known for its importance as migratory bottleneck for many species of birds and in particular for soaring raptors [46]. Experienced birdwatchers carried out daily observations near the radar to characterize species composition and migration traffic rate [47]. Radar measurements and direct observations of migrating birds took place daily during the study period, continuously between sunrise and sunset (UTC + 1), interrupted only occasionally by heavy precipitation. Radar tracks were generated using Hypatia-trackRadar, a dedicated software that uses a supervised approach that minimizes errors when estimating bird tracks from the radar echoes [48]. The program translates the echoes to a metric coordinate system and based on time and spatial position of the echoes, it automatically calculates multiple flight parameters, such as groundspeed, track length, and duration. In addition, the software allowed us to associate the echoes with the species observed by the birdwatchers. We filtered out tracks less than 100 m long and with a groundspeed of less than 5 m/s. Data regarding processed tracks are available in the DRYAD repository (doi: 10.5061/dryad.8gtht76q8).

**Figure 1:**
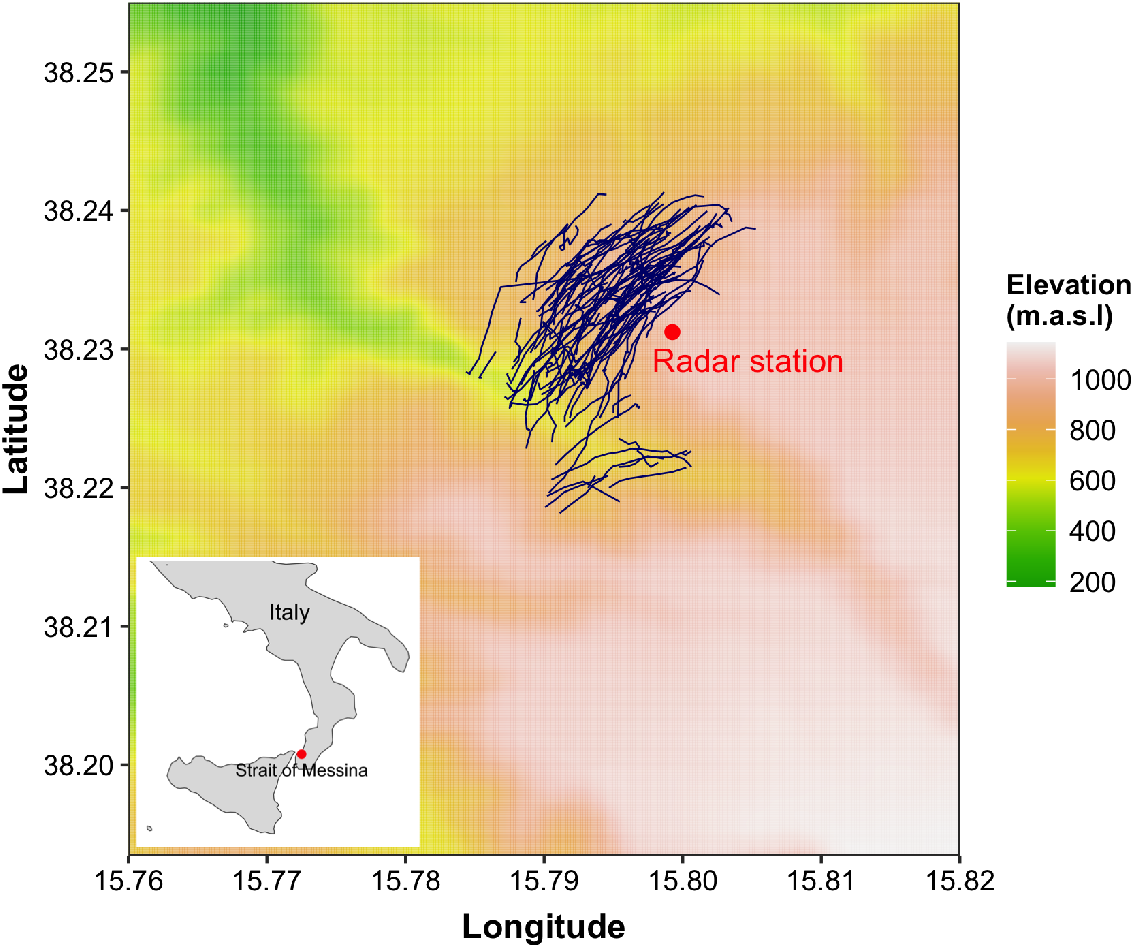
Map of the study area showing the position of the radar and observers (red dot). Radar tracks of one day (20/04/2017) are depicted in dark blue. These tracks were recorded in a day without fog. Inset figure shows the study location in Southern Italy, in close proximity with the Strait of Messina.

### Weather data

The presence of fog and low clouds was visually assessed by the radar operators and the birdwatchers. It was recorded (presence/absence) by assigning a presence value for each hour in which visibility was lower than 0.3 km for at least 15 consecutive min, disregarding isolated passing clouds (for details on the method, see [17]). Hourly U (eastward) and V (northward) components of the wind (10 m above ground) were downloaded from the ERA5 dataset of the European Centre for Medium-Range Weather Forecast (ECMWF) repository. ERA5 provides hourly estimates of a large number of atmospheric variables on a 30 km grid. We could not assess the exact height of the flying birds, hence we assumed that wind speed at 10 meters above the ground level was correlated to the wind speed at the height of birds’ flight [9, 49]. We used the wind components to calculate tailwind and crosswind speeds (hereafter TW and CW, respectively) relative to the mean seasonal (springtime) migration direction of the birds *θ_b_*, which is considered as their overall heading [33, 50]:

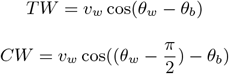

where *θ_w_* is the mean wind direction and *v_w_* is the wind speed. We noted that fog occurred only when the wind was blowing towards the sector that is between 90.92°and 179.22°. Thus, we filtered out all bird tracks recorded when the wind blew in directions beyond/outside this angular sector. In addition, we retained only tracks for which wind speeds were lower than 7.2 ms^-1^, since foggy conditions were almost always above this wind speed (Fig. S1).

### Movement parameters

To test if bird movement changed under different wind and fog conditions, we calculated several variables derived from their radar tracks, including:

1. Groundspeed: we calculated it as the simple ratio between distance covered and time passed between two consecutive points *v_g_*. We calculated the groundspeed component relative to the mean direction of migration *θ_b_* as a projected segment towards that specific direction *v_g_M*:

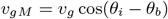

where *θ_i_* is the mean direction of the individual track.
2. Airspeed: we calculated it like [40], using the wind components crosswind CW and tailwind TW which were calculated in relation to the mean direction of migration *θ_b_*:

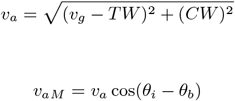

where *v_a_* is the airspeed in relation to TW and CW experienced by the buzzards, and *v_a_M* is the *v_a_M* component relative to the mean direction of migration *θ_b_*.
3. Sideways speed: we calculated the bird’s lateral speed relative to *θ_b_* with the following formula:

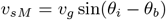

All angles are expressed in radians. We calculated the mean migration direction *θ_b_* as the circular mean of all tracks collected from buzzards that passed through the radar’s detection area in fair weather only. Furthermore, we used hourly means of bird movement parameters to match the temporal resolution of the wind measurements. In addition, we filtered out tracks that deviated more than 100°from the *θ_b_* (either to the left or to the right of this angle) to exclude the movement of local and non-migrating birds.

### Statistical analysis

We used the Watson’s U^2^ test (Watson’s Two-Sample Test of Homogeneity) to compare between the two sets of track directions, under fog and without fog [51]. We modelled hourly means of groundspeed *v_g_M*, airspeed *v_a_M*, and sideways speed *v_s_M* as functions of the tailwind and the crosswind components of the wind (continuous variables) and fog presence (binomial). Wind data is at the same temporal resolution (hourly values) to avoid eventual pseudo-replication (having many tracks recorded in a certain hour that are matched with a single wind parameter value). We used linear mixed effect models (LMMs), with ordinal date as an intercept random effect. We found the optimal structure of the fixed components of the model and ran the models with different combinations of the fixed effect terms (tailwind, crosswind and fog presence) in the global model. We used the log-transformed count of tracks per hour as weights in the model. To evaluate model fit, we used the Akaike Information Criterion (AIC; [52]), by applying the function *dredge* from the `MuMIn` package in R [53, 54]. Following inspection of model residuals and considering the dispersion of the data using the `DHARMa` package [55], we chose linear mixed models as the most appropriate [56]. Statistical analyses were performed in R 4.1.0 [54] using packages `glmmTMB` [57] and `circular` [58]. Plots and table of the models were produced using the package `sjPlot` [59] and `ggplot2` [60].

## Results

We recorded a total of 60,552 radar tracks; of these, 2,885 (4.8%) were recorded during foggy conditions (Fig. 2, see also Fig. S2 for the number of tracks per day in fog and fair-weather conditions). After filtering out tracks that did not meet the wind speed and direction criteria (as explained in “Methods”), we retained 28,553 tracks. The mean flight direction of birds differed significantly between birds that flew under clear skies (96% of the tracks) and fog (4% of the tracks) conditions (Watson’s U^2^ test: test-statistic = 2.99, p < 0.001), with a mean direction of 57.3°under clear skies and 80.7°under foggy conditions (Fig. 2). Mean hourly values for groundspeed, airspeed and sideways speed were computed for a total of 270 hours of radar operation. The selected statistical models with groundspeed and sideways speed relative to the mean migration direction as dependent variables did not include tailwind and its interaction with fog condition, while the model selected for explaining bird airspeed retained tailwind but not its interaction with fog presence (Table 1). Soaring migrants flying in foggy and non-foggy conditions over the study area had, in general, similar groundspeeds, but they differed in their response to crosswind speed (Fig. 3). Increased crosswind speed under clear skies induced a decrease of groundspeed, while groundspeed increased with crosswind speed under foggy conditions (Table 1; Fig. 3). It should be noted that due to the filtering of the tracks, the crosswind component contains only positive values, indicating a wind blowing towards the bird’s right-hand side (Fig. 3-5). Under clear skies and foggy conditions, birds reduced their airspeed with increasing tailwind speed, as expected (Table 1; Fig. 4A). With increasing crosswinds, buzzards flying under clear skies decreased their airspeed while those flying under fog increased their airspeed (Fig. 4B). Sideways speed differed between buzzards flying in fog versus clear air (Table 1; Fig. 5A). In foggy conditions, sideways speed increased towards the right side of the migration goal (positive values), in the inland direction. Birds travelling under clear skies had a negative sideways speed, meaning that they moved more towards the coast (Table 1; Fig. 5A). The general response of the birds to crosswind was slightly different; under clear skies they tended to direct their flight towards inland areas under weak crosswinds and towards the coast with increasing crosswind speed (Table 1; Fig. 5B). When crosswinds were weak, birds flew towards the general direction of the presumed migration goal (dashed line in Figure 5), but under the same crosswind conditions, bird that flew in the fog over-compensated for wind drift by traveling towards the coast. With increasing crosswind speed, the birds eventually compensated for the lateral drift by flying towards the presumed migration goal (Fig. 5B). For birds travelling in foggy condition, the sideways speed varied too much to establish a clear response, but their response clearly differed from that of birds flying in clear weather.

**Figure 2:**
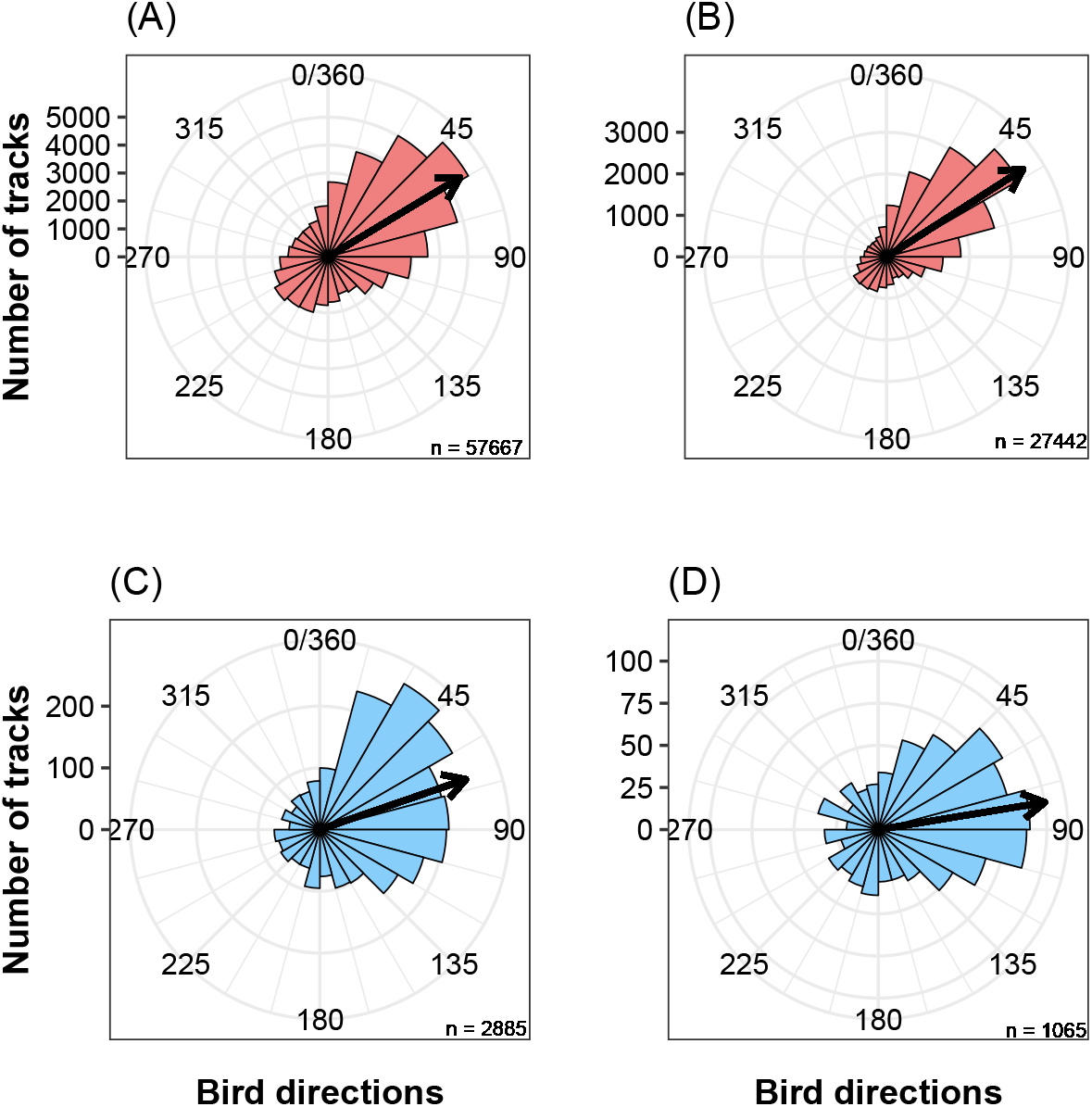
Circular distribution of (A) all tracks recorded under clear skies, (B) tracks recorded under clear skies in selected wind conditions that match those recorded during fog events, (C) tracks recorded under fog conditions, and (D) tracks recorded under fog conditions in selected wind conditions. Black arrows are mean circular directions.

**Figure 3:**
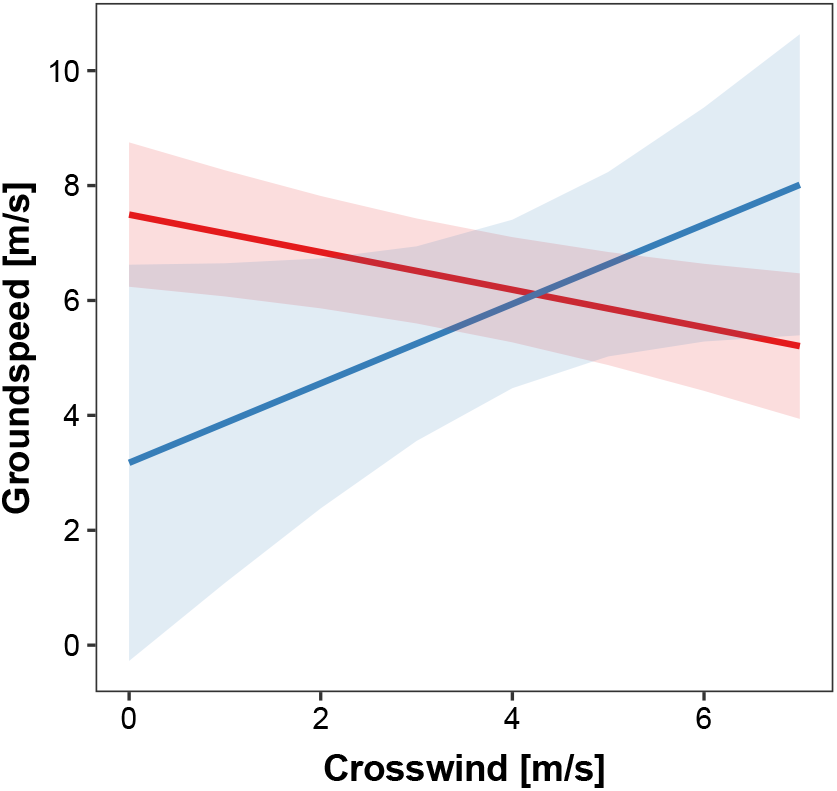
The effects of crosswind on groundspeed of Honey Buzzards migrating through fog (blue) and under clear skies (red). Regression slopes and 95% C.I. of the groundspeed in relation to crosswind in foggy conditions (blue) and clear weather (red). Crosswinds all have positive values since they only represent wind blowing from the left to the right side of the average migration direction.

**Figure 4:**
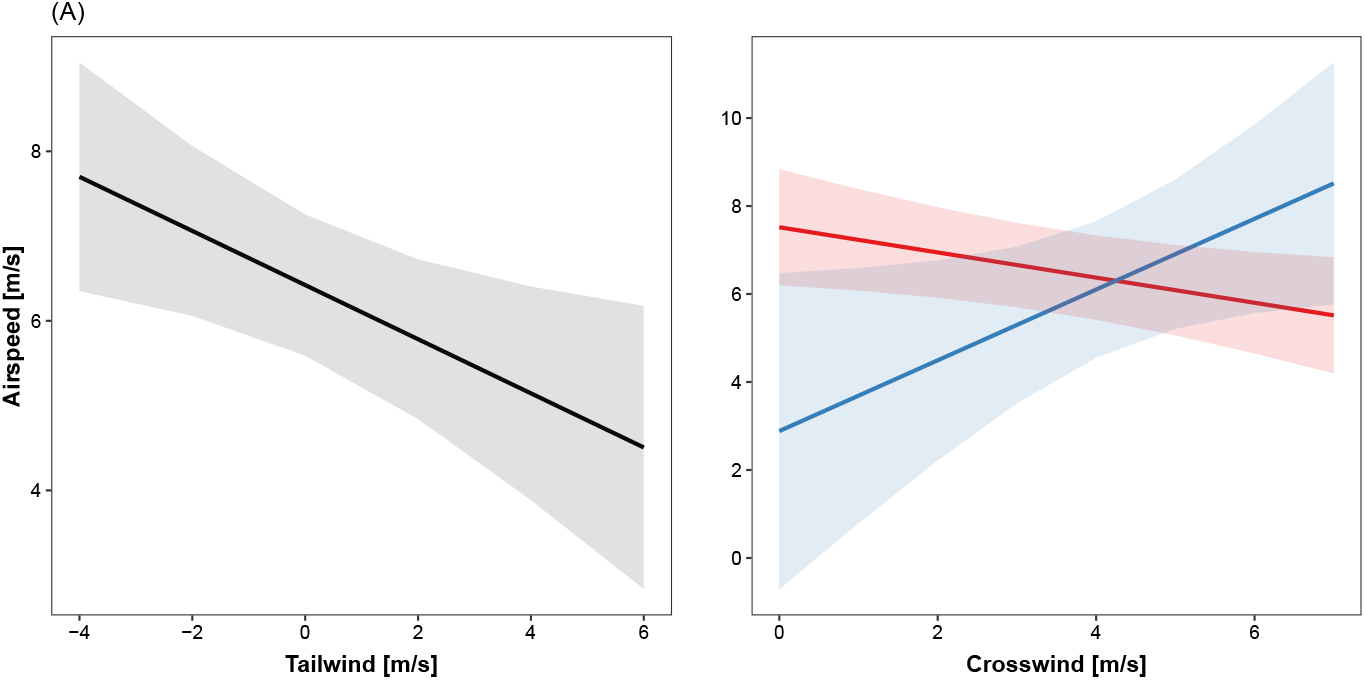
The effects of crosswind and tailwind speed on airspeed of Honey Buzzards migrating through fog or under clear skies. (A) Regression slope and 95% C.I. of the airspeed in relation to tailwind. (B) Regression slopes and 95% C.I. of the airspeed in relation to crosswind in foggy conditions (blue) and clear weather (red). Crosswinds all have positive values since they only represent wind blowing from the left to the right side of the average migration direction.

**Figure 5:**
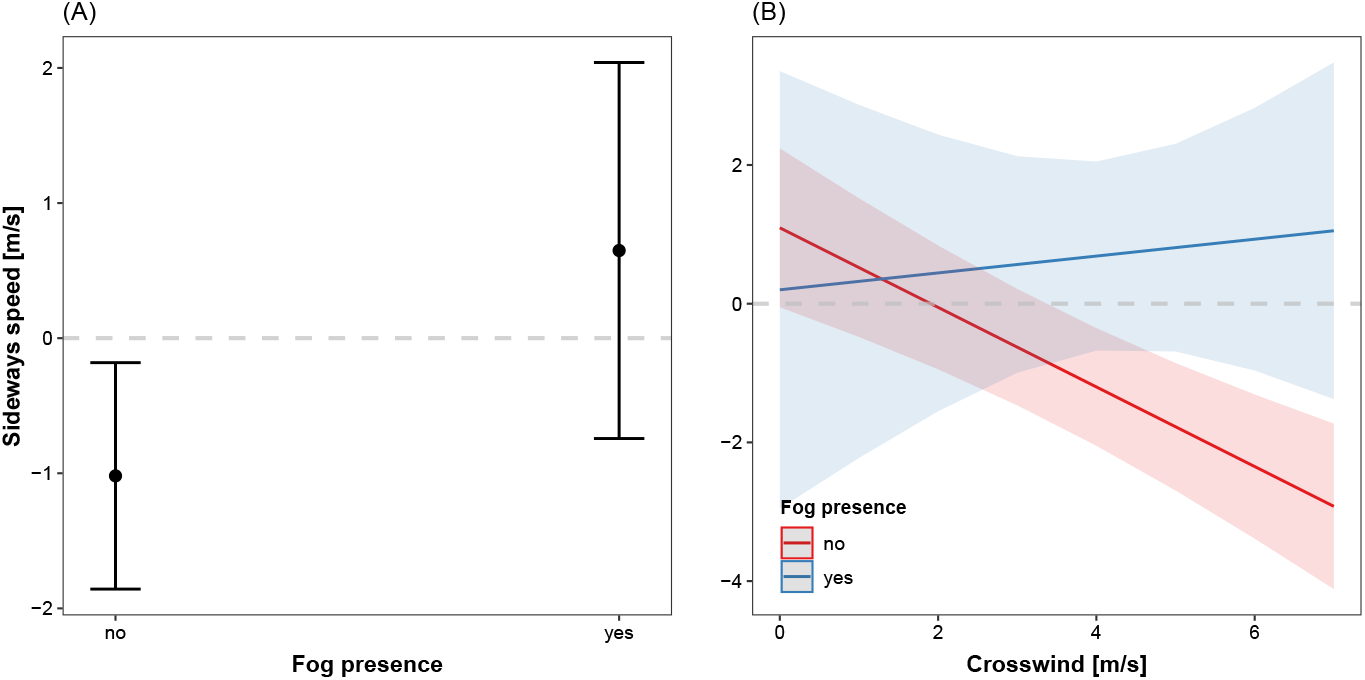
The effects of crosswind on bird sideways speed. (A) Sideways speed mean and 95% C.I. (whiskers) for tracks in foggy and clear weather conditions. (B) Regression slopes and 95% C.I. of the sideways speed in relation to the crosswind component through fog or under clear skies. The dashed line marks the y-intercept at 0 m/s sideways speed, representing the presumable direction of flight towards the migration goal. Crosswinds have only positive values since they only represent the wind blowing from the left to the right side of the average migration direction.

**Table 1:**
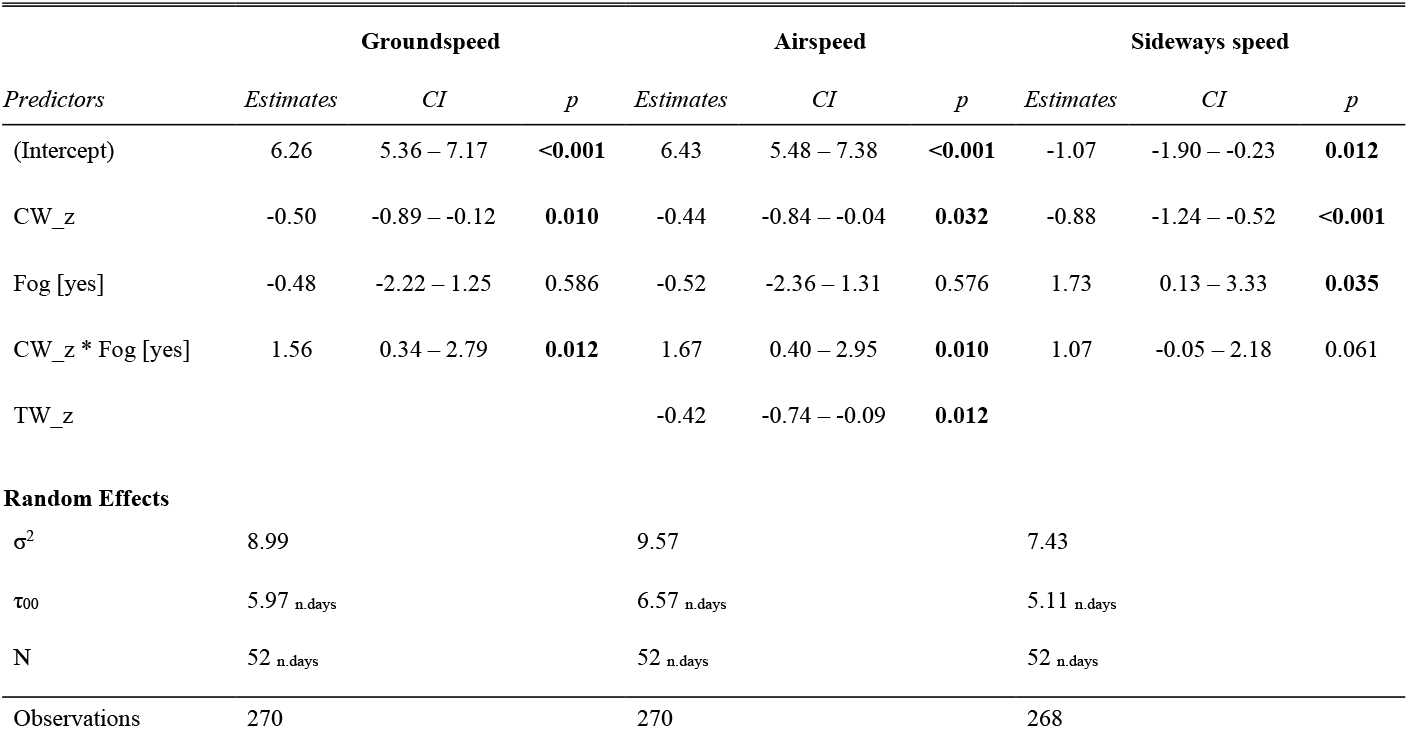
The selected linear mixed models reporting the effects of wind and fog on bird average hourly speeds. Continuous predictors are scaled (z-transformed). Significant p-values are in bold.

## Discussion

Our findings demonstrate that fog affects bird orientation and the modulation of flight speed in relation to wind during migration. We found that the birds’ flight directions were more scattered under foggy conditions and that the mean migration direction changed when fog was present. As described in an earlier study, soaring migrants’ densities are drastically (95 %) reduced in fog [17], and the present study uncovers the reasons for this avoidance, suggesting that bird orientation can be disrupted and the cost of transport could be elevated such that the migration trip may become more risky and costly. For example, birds try to bypass fog by displacing themselves in an unpredictable manner or to distant places [15, 19], or worse, they might hit a natural or artificial obstacle obscured by fog. To avoid this risk, birds flying in fog often land and wait until the fog clears to resume their migration [16, 21]. Fog may make spotting key landmarks necessary for maintaining a certain route direction diffcult, consequently influencing bird behaviour and movement. We found that soaring birds flying under both clear skies and foggy conditions modulated their ground and air speed in relation to tail-and headwind as predicted by optimal migration flight theory [34, 35], whereas bird response to the crosswind component of the wind was different under foggy conditions. The increase in airspeed, and consequently groundspeed, with increasing crosswinds could help birds to get out of areas covered by fog. Yet, such behaviour may expose birds to higher risk of collision with structures obscured by the fog, such as buildings, communication towers, power lines and wind turbines [21, 61, 62]. Furthermore, changes in bird sideways speed in foggy conditions could result in more scattered tracks that diverge from the intended migration goal, as can be expected under limited visibility conditions in general [16, 19, 21] and observed in this study. Soaring migrants that faced increasing crosswinds (from sea to land, as in Fig. 5B) under clear skies tended to fly towards land, but during foggy events, sideways speed was very variable and this can be reflected by the scattered mean track directions around the migration goal vector (Fig. 2C-D). It seems that the birds were ranging between under-compensating for lateral drift to over-compensating as the crosswind component increased in clear weather and this response differed under foggy conditions. These findings are the first illustration of how fog influences the flight properties of soaring migrants. The lack of data on the spatial and temporal properties of fog in meteorological databases limits broad-scale analysis of the effects of fog on migrating animals; thus, only a handful of small-scale studies have been conducted so far to study these phenomena [17, 18]). Some of these studies investigated orientation behaviour under fog, including a study that found that fog prevented Little Penguins (*Eudyptula minor*) from crossing land to reach their nests walking, probably affecting their orientation and delaying their nest attendance [14]. Other studies, mostly anecdotal, suggested that fog also affected migrating dragonflies [15], migrating storks [16] and vultures [19]. In nocturnal species, fog amplifies the glow of artificial lights and influences the orientation of birds [21, 63] and insects [5]. As already proposed by Panuccio et al. [17], we believe that the consequences of flying in fog must be considered when estimating bird collision risks in human-made structures in areas where fog is common. The scatter of flight directions under foggy conditions that we documented could be genuine but might also be explained by our inability to properly define the spatial extent and the intensity of the fog around the radar station. It is possible that some birds were outside the fog when tracked (or perhaps above it), and some experienced different visibility of landmarks when flying in the fog due to variations in fog intensity. We occasionally saw some of the buzzards during the fog events near the radar site, and noticed that the birds dramatically decreased their flight elevation and flew very close to the ground, perhaps to see landmarks. The possibility to measure altitude along with changes in the birds’ horizontal position would be of great use to explain flight behaviour in low visibility conditions in future studies. We note that radar technology provides a valuable means for collecting data and quantifying behavioural changes in birds flying in fog, as direct observations of birds in these conditions are nearly impossible [5]. Our findings show that birds flying in fog do not adjust their flight behaviour as well as they do in good visibility conditions, with possible consequences that could lower bird survival. Finally, our findings represent the first empirical evidence of the variation of flight behaviour under foggy conditions in migratory birds, highlighting how birds may cope with the risks of flying in low visibility conditions.

## Acknowledgements

Data collection was supported by Terna Rete Italia S.p.A. We thank Ornis italica which supported our fieldwork. We thank all the people involved in the field as radar operators and as birdwatchers. MP was partially funded by COST-European Co-operation in Science and Technology through the Action ES1305 European Network for the Radar Surveillance of Animal Movement (ENRAM). We would like to thank G. Bohrer and E. Kirsch for their useful comments that substantially improved our work, and S. Turjeman for the English revisions.

