## Supplementary material for "Groping in the fog: soaring migrants exhibit wider scatter in flight directions and respond differently to wind under low visibility conditions": Fig. S1 Fig. S2 Fig. S3

**Electronic Supplementary Materials of the article titled:**

### "Groping in the fog: soaring migrants exhibit wider scatter and do not properly adjust their flight in response to wind under low visibility conditions"

Paolo Becciu[1,2,3]*, Michele Panuccio[2,‡], Giacomo Dell'Omo[2] and Nir Sapir[3]

[1] Department of Ecology and Evolution, University of Lausanne, Lausanne, Switzerland.

[2] Ornis italica, Roma, Italy.

[3] Department of Evolutionary and Environmental Biology and Institute of Evolution, University of Haifa, Israel.

[‡] Deceased.

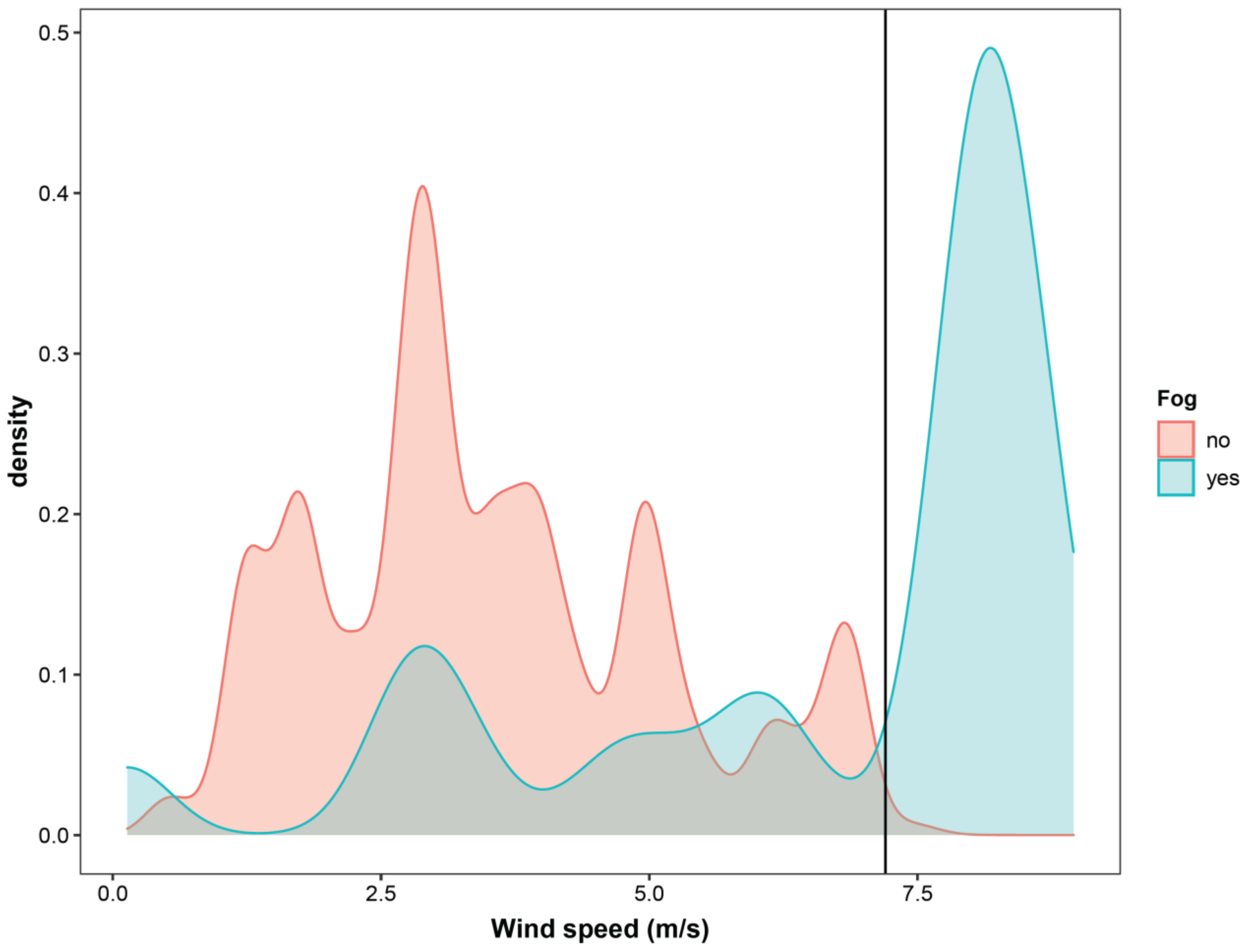


**Figure S1** | Distribution of wind speeds experienced by the soaring migrants flying under clear skies or foggy conditions during the study period. The portion on the right side of the graph is representing the range of data excluded in the analyses (wind speed > 7.2 m/s).

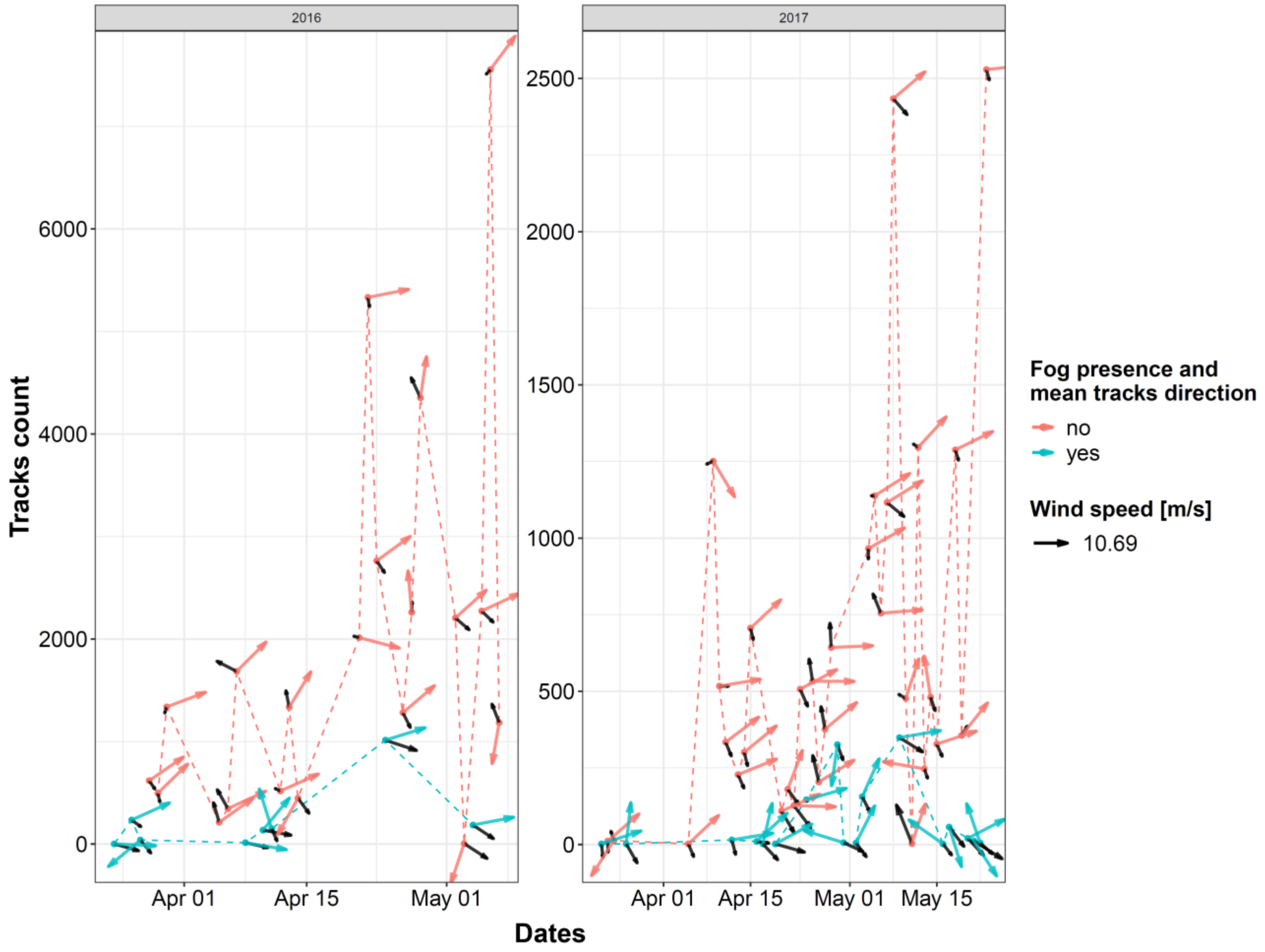


**Figure S2** | Number of radar tracks recorded per day in 2016 and 2017. Tracks are grouped by fog condition and mean daily directions which are expressed by the coloured vectors. Black arrows are referring to the mean wind speed (vector length) and direction.

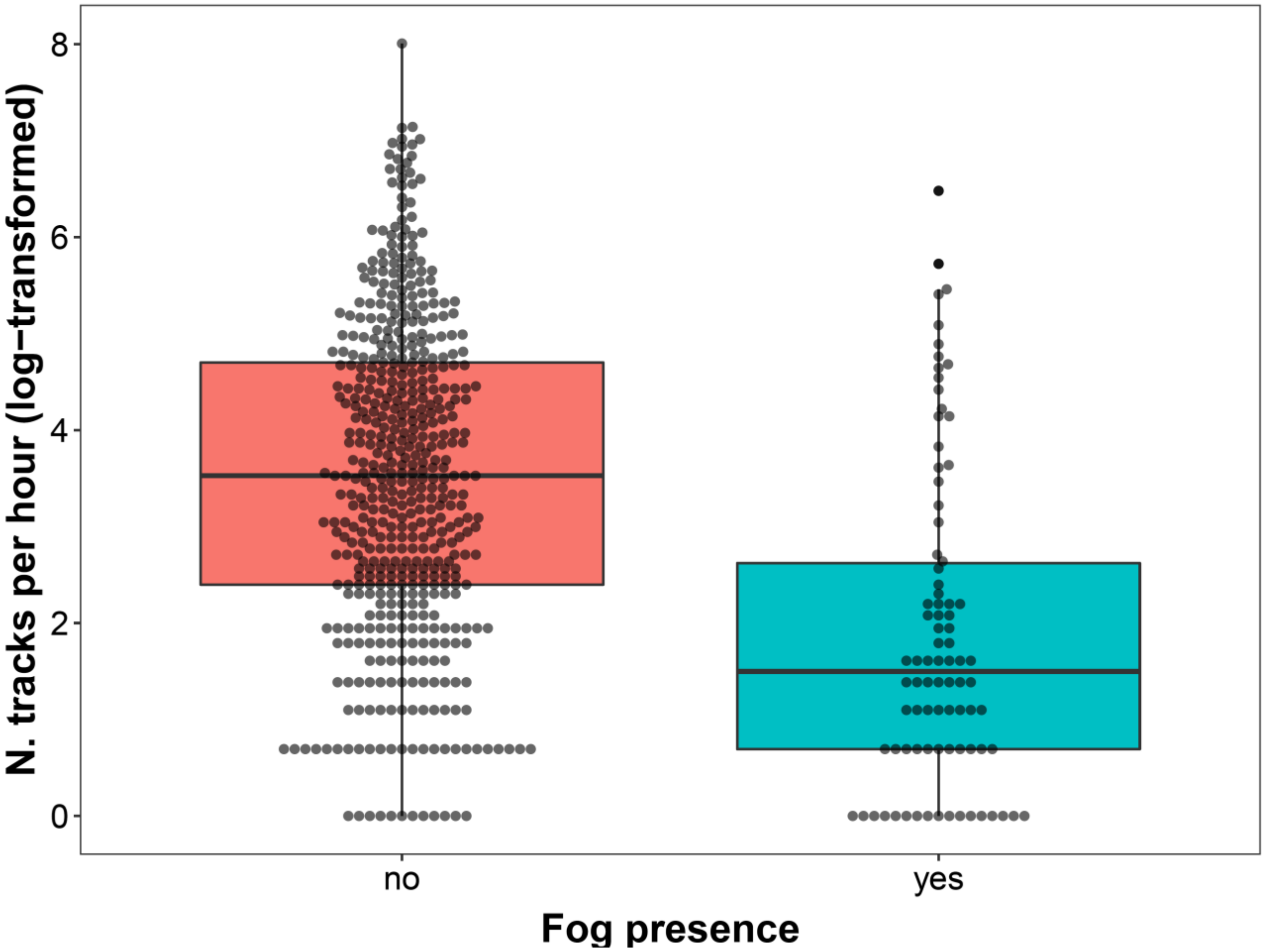


**Figure S3** | Hourly tracks recorded under fog and clear skies. Data includes all the collected tracks, averaged per hour. Black line represents the median and box is the interquartile range. The upper whisker is the maximum value within 1.5 times the interquartile range over the 75th percentile. The lower whisker is the minimum value within 1.5 times the interquartile range under the 25th percentile.
